# SynergyAge: a curated database for synergistic and antagonistic interactions of longevity-associated genes

**DOI:** 10.1101/2020.04.22.054767

**Authors:** Gabriela Bunu, Dmitri Toren, Catalin-Florentin Ion, Diogo Barardo, Larisa Sârghie, Laurentiu Gabriel Grigore, João Pedro de Magalhães, Vadim E. Fraifeld, Robi Tacutu

## Abstract

Interventional studies on genetic modulators of longevity have significantly changed gerontology. While available lifespan data is continually accumulating, further understanding of the aging process is still limited by the poor understanding of epistasis and of the non-linear interactions between multiple longevity-associated genes. Unfortunately, based on observations so far, there is no simple method to predict the cumulative impact of genes on lifespan. As a step towards applying predictive methods, but also to provide information for a guided design of epistasis lifespan experiments, we developed SynergyAge - a database containing genetic and lifespan data for animal models obtained through multiple longevity-modulating interventions. The studies included in SynergyAge focus on the lifespan of animal strains which are modified by at least two genetic interventions, with single gene mutants included as reference. SynergyAge, which is publicly available at www.synergyage.info, provides an easy to use web-platform for browsing, searching and filtering through the data, as well as a network-based interactive module for visualization and analysis.

Database URL: http://www.synergyage.info/

**BACKGROUND & SUMMARY:** The aging process can be genetically modulated. This has been previously shown in many studies in which average lifespan, and in some cases even maximum lifespan, has been modified by genetic interventions. It is for example possible to have genetic mutants with an increased lifespan, up to ten times higher compared to wild type in C. elegans ^1^, and up to 150% and 46% in *D. melanogaster* and *M. musculus,* respectively ^2,3^. Up until now, at least 2,205 genes, whose mutations, downregulation or overexpression results in a long- or short-lived phenotype, have been identified in model organisms. A comprehensive list with these longevity-associated genes (LAGs), including more detailed information about lifespan experiments, can be found in the GenAge database ^4^. This type and amount of data have made it possible for higher level analyses to be performed ^5–7^, and the collection of LAGs in public repositories has significantly pushed biogerontology towards more integrative approaches to study longevity. One important aspect observed is that many LAGs seem to act in a cooperative manner ^8–10^ and are not independent regulators of lifespan. In fact, in most cases when combining two or more genetic interventions, the effect is rarely additive, as genes are generally epistatic and interact in nonlinear ways ^11,12^. While in most cases combined interventions seem to have lower than expected results in how much they extend lifespan, there are also a minority of cases where genes act synergistically ^13,14^. Even so, the much more common case is that of studies where partially dependent gene interactions are revealed, making it even more important to understand and predict genetic dependencies.

Unfortunately, data on epistasis is much harder to obtain through wide-screen experimental studies, which has been for example the case for the discovery of most LAG interventions in worms. The main impediment comes from the combinatorial explosion of multiple gene groups for which lifespan assays would need to be measured in a “blind” search, through wet-lab experiments. Instead, it would be more efficient to use existing epistasis data to explore predicted synergies in guided lifespan experiments. Luckily, an accumulating number of papers has been published in the last two decades with reported lifespans for double mutants and in some cases even triple or quadruple mutants. As such, it has been now possible for us: (i) to collect the data from existing studies containing lifespan records for strains that have multiple genes modulated, and (ii) to create an intuitive, network-based tool, which allows users to explore in a fast, visual and interactive way the lifespan relationships between these strains.

Here, we present SynergyAge, a database containing manually curated data, extracted from experimental studies, regarding gene combinations that affect lifespan. With the creation of SynergyAge, we aim to encourage the investigation of the cumulative effects of different gene interventions on lifespan, by providing the scientific community with a “one-stop” web platform to access, compare and analyze lifespan synergisms or antagonisms. This resource is of particular interest in designing wet-lab experiments in which multi-genetic lifespan interventions are needed. SynergyAge (http://www.synergyage.info) is publicly available and contains data from three animal model organisms: *Caenorhabditis elegans*, *Drosophila melanogaster* and *Mus musculus*.

## METHODS

## Notations and definitions

In order to simplify the language of the article (and the website interface), two notations have been adopted: 1) animal strains upon which genetic interventions, including mutations, knockout, overexpression or RNA interference, were carried out, are loosely referred to as “mutants” (as opposed to wild type strains); 2) animal strains with a number (n) of molecular interventions performed in different genes are called n-mutants (e.g. a double mutant is also called 2-mutant while a triple mutant is referred to as a 3-mutant).

### Data sources and data curation

Only manually curated data from scientific articles were used for the content of SynergyAge. Experiments were included in the database if: 1) lifespan assay was carried out for an n-mutant and was compared to the lifespan of at least one strain from which the mutant was incrementally constructed - i.e. an (n-1)-mutant, and 2) at least one of the mutants in the experiment showed a lifespan increase greater than 10% compared to the wild type. In exceptional cases, when the lifespan value of the wild type was not assessed in the same study, experiments were still included in the database if one of the genetic interventions was equivocally shown to modulate lifespan in prior studies. It should be mentioned that due to the uncertainty in target specificity, mutants with drug-modulated longevity have not been considered in SynergyAge.

Information about experimental conditions was manually extracted from the included studies. Gene-centric data was automatically annotated from Entrez Gene ^15^ and organism-specific databases Wormbase ^16^, Flybase ^17^ and MGI ^18^.

In the cases where the survival curves in the paper hint towards animals suffering (and dying) from any specific pathologies (e.g. sudden deaths across cohort, severe differences in the control group compared to the literature), a more thorough curation was done and the articles were excluded if the cohorts were not representative for “normal” aging. In particular, for short-lived mutants, we generally checked if the authors reported signs of premature aging (e.g. graying of hair in mice, incremental slowness in feeding patterns or movement in worms, etc) being observed as a slow natural decline, and if no observations were made towards any specific pathologies.

### Evaluation of epistasis: synergism, antagonism and monotony

When new builds are created for SynergyAge, a script automatically categorizes the curated gene combinations, adding annotations about the epistatic interactions of each n-mutant. If data is available for all intermediary mutants, then “full” synergism is evaluated. If intermediary data is missing, the partially known monotony is computed instead.

For 2-mutants, the only category for which “full” synergism can be evaluated, epistatic interactions have been classified as synergistic, additive, almost additive, dependent or antagonistic, based on the difference between the effect of the combination and the sum of the effects of individual genetic interventions. Briefly, given the lifespan (LS) for wild type (WT), gene 1 mutant (G1), gene 2 mutant (G2) and the double mutant (G1,G2), we defined the effects on lifespan, compared to wild type, for each model as follows:

ΔG1 = (LS(G1) – LS(WT)) * 100 / LS(WT)

ΔG2 = (LS(G2) – LS(WT)) * 100 / LS(WT)

Δ(G1,G2) = (LS(G1,G2) – LS(WT)) * 100 / LS(WT)

Additionally, for epistatic classifications only combinations where all single gene interventions have the same direction effects were evaluated: I) ΔG1>0, ΔG2>0 (both interventions have positive effects on lifespan) and II) ΔG1<0, ΔG2<0 (both interventions have negative effects on lifespan). Below are the rules for all annotated categories:

- Fully synergistic interaction: |Δ(G1,G2)| > |ΔG1| + |ΔG2|
- Additive interaction: |Δ(G1,G2)| = |ΔG1| + |ΔG2|
- ‘Almost’ additive interaction: max(|ΔG1|, |ΔG2|) < |Δ(G1,G2)| < |ΔG1|+ |ΔG2|
- Antagonistic, dependent interaction: min(|ΔG1|, |ΔG2|) <= |Δ(G1,G2)| <= max(|ΔG1|, |ΔG2|)
- Fully antagonistic: |Δ(G1,G2)| < min(|ΔG1|, |ΔG2|)

If the lifespan of the double mutants is changed into the opposite direction compared to the lifespan of single mutants, the relationship is also considered antagonistic.

For n-mutants that do not have all paths known, a monotony evaluation is carried out. Briefly, this is performed by checking that on each known path from wild type to the n-mutant, every intervention (k) increases the lifespan of the previously constructed mutant (combining k-1 gene interventions). If at least one non-monotonic path is found, then it is considered that the combination of n genes contains dependency relationships.

### Data visualization and interface

All networks used for the browsing of organism-specific data were created using Cytoscape 3.5.1 and are displayed on the website using Cytoscape.js ^19^. The change in average lifespan displayed in the interface represents the interval defined by the minimum and maximum effects obtained for a gene intervention or a combination of gene interventions in all the experiments recorded in the database.

If the lifespan of the wild type was not assessed in the same experiment as the mutant, the displayed upper and lower limits of the interval are calculated by comparing the lifespan of the mutant to the maximum and the minimum lifespan reported in all studies, for the wild type of that species.

To create a unique ordered list, the names of n-mutants are formed by concatenating the names of the mutated genes in alphabetical order (i.e., daf-16;daf-2 and daf-2;daf-16 are both displayed as daf-16;daf-2). This notation was chosen in order to aggregate the reported data from studies in which the order of the performed genetic interventions varied.

## DATA RECORDS

### Lifespan synergism and antagonism

In order to clearly separate between different types of epistatic interactions, four types of relationships between genes have been defined based on the difference between the effect of the combination and the sum of effects of individual genetic interventions. In Fig. 1, these relationships are exemplified for 2-mutants: 1) for “synergistic” genes, the effect of the gene combination is greater than the sum of the individual effects; 2) genes with “additive” and “almost additive” effects are those for which the effect of the combination is greater than each of the two individual effects but weaker than their sum (the case for “additive” is mostly theoretical and happens when the effect of the combination is exactly the same as the sum); 3) for “dependent” genes, the effect of the gene combination is a weighted average of the individual effects, suggesting that the greater individual effect requires the other gene to be unaltered, and 4) genes with “antagonistic” effects are those for which the effect of the combination is worse than any of the individual effects. For a wider examination of epistasis with regards to n-mutants (n>2), please see the Discussion subsection.

**Fig. 1.**
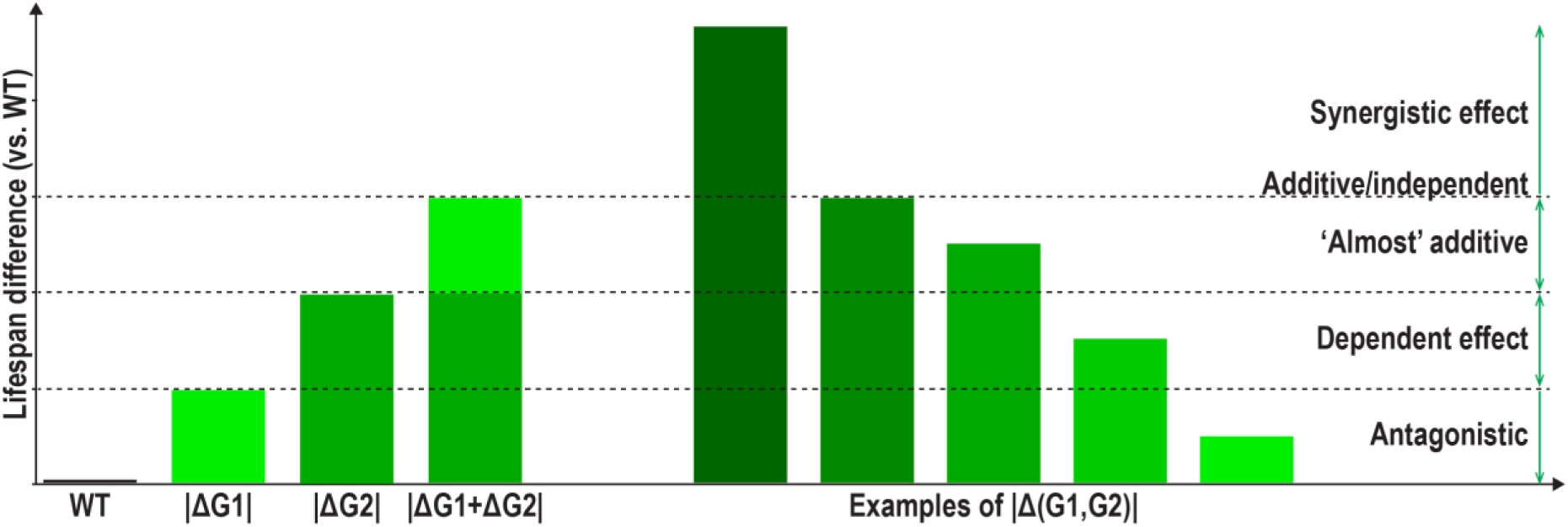
Lifespan epistasis categories defined for combinations of two genetic interventions. Depicted are synergistic, additive, almost additive, dependent and antagonistic relationships, categorized based on the difference between the effect of the combination and the sum of the effects of individual genetic interventions. By these definitions, we are considering that individual gene interventions resulting in changes with opposite lifespan modulations cannot be directly assessed for epistasis.

### Database Content

SynergyAge contains longevity data characterizing mutant strains obtained through a series of genetic interventions for which lifespan assays have been performed. These studies include animals with at least two genetic interventions, whose role is to either inhibit (loss-of-function mutations, knockout or RNA interference) or activate (gain-of-function mutations or overexpression) gene function. For the purpose of comparing the effect on lifespan that multiple genes vs single genes have, the database also contains information about the lifespan of single gene mutants that have been used intermediately to construct the multiple gene mutants. Besides lifespan values and a summarized description of the reported results, SynergyAge hosts information about the experimental conditions that might have affected lifespan, including the temperature at which the experiment took place, and the diet that was administered to the animals. Additionally, for each of the n genes modulated in a combined intervention, the database provides the user with a panel that contains direct links to general gene/protein resources, like Entrez Gene, Ensembl, Uniprot as well as to organism-specific databases, like Wormbase, Flybase or MGI. In these gene-centric panels, direct links to GenAge entries are also provided for quick access to longevity-related information, if those genes are already known LAGs.

In some cases, gene combinations may be annotated with more than one epistatic relationship due to different results being reported in the literature. For such entries, we aim to be inclusive, conveying the entire information to the user.

It should be mentioned that the majority of studies present important lifespan data without fully considering the epistatic aspects - generally without assaying all the intermediary mutant strains. However, considering the current *status quo* of the field, this type of “incomplete” data is much needed to increase our understanding of the genetic epistasis of longevity, and thus it is also included in SynergyAge.

### Statistical description

Currently the database contains almost 7000 lifespan records for more than 6200 genetic mutants. Of these more than 3600 mutants have at least 2 mutations, with the rest providing comparative information as single mutants. In total, 794 unique genes have been identified as part of multi-genetic interventions, in three model animal organisms: *Caenorhabditis elegans*, *Drosophila melanogaster* and *Mus musculus*(Table 1).

**Table 1.** Number of records in the SynergyAge database

|  | *C. elegans* | *D. melanogaster* | *M. musculus* | Total |
| --- | --- | --- | --- | --- |
| Lifespan values | 6656 | 185 | 147 | 6988 |
| Gene combinations | 1770 | 27 | 3 | 1833 |
| Genes in total | 714 | 36 | 44 | 794 |
| Articles | 144 | 20 | 29 | 193 |

As expected, the large majority of records hosted in SynergyAge is for *C. elegans*, because the amount of lifespan data available in the literature for this species is significantly higher than for the other two organisms. However, data for *Drosophila melanogaster and Mus musculus,* is beginning to accumulate and as such, dedicated sections for flies and mice have been created.

Out of the 794 genes included in SynergyAge, 353 have already been associated with the modulation of lifespan through genetic experiments, and are reported in GenAge as LAGs ^4^, while others are either new LAGs or work in conjunction with known LAGs. Of previously reported LAGs, manipulations of 220 genes resulted in a lifespan increase and 108 in a lifespan decrease of at least 10%, compared to the wild type.

Currently, a large part of the n-mutants with n>1 is represented by double mutants (N=1254), with a smaller number of studies in which triple mutants (N=491) or quadruple mutants (N=84) have been assayed. As defined above, double mutants can be easily associated with an epistasis category (see Fig. 1), depending in each case on the availability of data about the two single mutants, and in such cases this annotation is done automatically. For the triple and quadruple mutants, most of which include genetic interventions either in the (daf-2, daf-16) or in the (fem-23, fer-15) tuples, very few of the intermediate mutants have been evaluated and thus synergism cannot be fully evaluated. Moreover, even for the double mutants, part of the studies present lifespan data only for one of the single mutants, making epistasis prediction only partial. In Table 2, a statistics summary is presented for the epistasis combinations among the double mutants with complete data.

**Table 2.** Statistics of epistatic relationships for all lifespan-modulating interventions combining the effect of two genes

|  | *C. elegans* | *D. melanogaster* | *M. musculus* | Total |
| --- | --- | --- | --- | --- |
| Long-lived phenotype |  |  |  |  |
| Synergistic | 190 | 2 | 0 | 192 |
| Additive | 2 | 0 | 0 | 2 |
| “Almost” additive | 85 | 5 | 0 | 90 |
| Dependent | 206 | 4 | 0 | 210 |
| Antagonistic | 50 | 6 | 0 | 56 |
| Short-lived phenotype |  |  |  |  |
| Synergistic | 9 | 3 | 3 | 15 |
| “Almost” additive | 23 | 3 | 4 | 30 |
| Dependent | 29 | 8 | 9 | 46 |
| Antagonistic | 42 | 3 | 0 | 45 |

### Discussion

The SynergyAge database is a repository of lifespans for genetic mutant strains and of inferred epistatic relationships between genes that were modified in order to obtain these long-and short-lived mutants. It should be mentioned, that in order to avoid potential divergences between the mechanisms of cellular aging and those of organismal aging, as well as potential divergences between chronological and replicative senescence, the database focuses only on animal model organisms, and hence *Saccharomyces Cerevisiae* was not considered in this project.

As seen above, defining the epistatic relationship for double mutants can be straightforward, provided that lifespan values are known for the wild type, single mutants and double mutant, kept under the same conditions. Similarly, epistasis categories (synergistic, additive, almost additive, dependent and antagonistic) could also be used for higher-level n-mutants (n>2), however, the inference rules need to be more complex, accounting for all the intermediary k-mutants (for any k<n).

#### Evaluation of “full” synergism

If we consider, for example, an n-mutant to be the result of molecular interventions in n genes, that act fully synergistic with regards to longevity, then it would be required that the lifespan of the n-mutant is greater than the sum of lifespans for any two intermediary k-mutant and (n-k)-mutant, where the subsets of k and n-k genes do not overlap (see Fig. 2A). An even stricter meaning of synergism would require that if n genes are synergistic, then any gene subsets have also to be synergistic amongst themselves (i.e., the statement above needs to be true for any m<n, where the m genes are randomly selected from the full gene set). This definition would however exclude combinations of genes that do not display synergism at a lower level (each pair of genes for example might not be synergistic), but that have a synergistic effect at a higher level (as an n-mutant). While choosing between the two definitions remains somewhat arbitrary, the first definition is more flexible and inclusive, which should be considered in light of the relatively limited number of synergistic interventions reported so far.

**Fig. 2.**
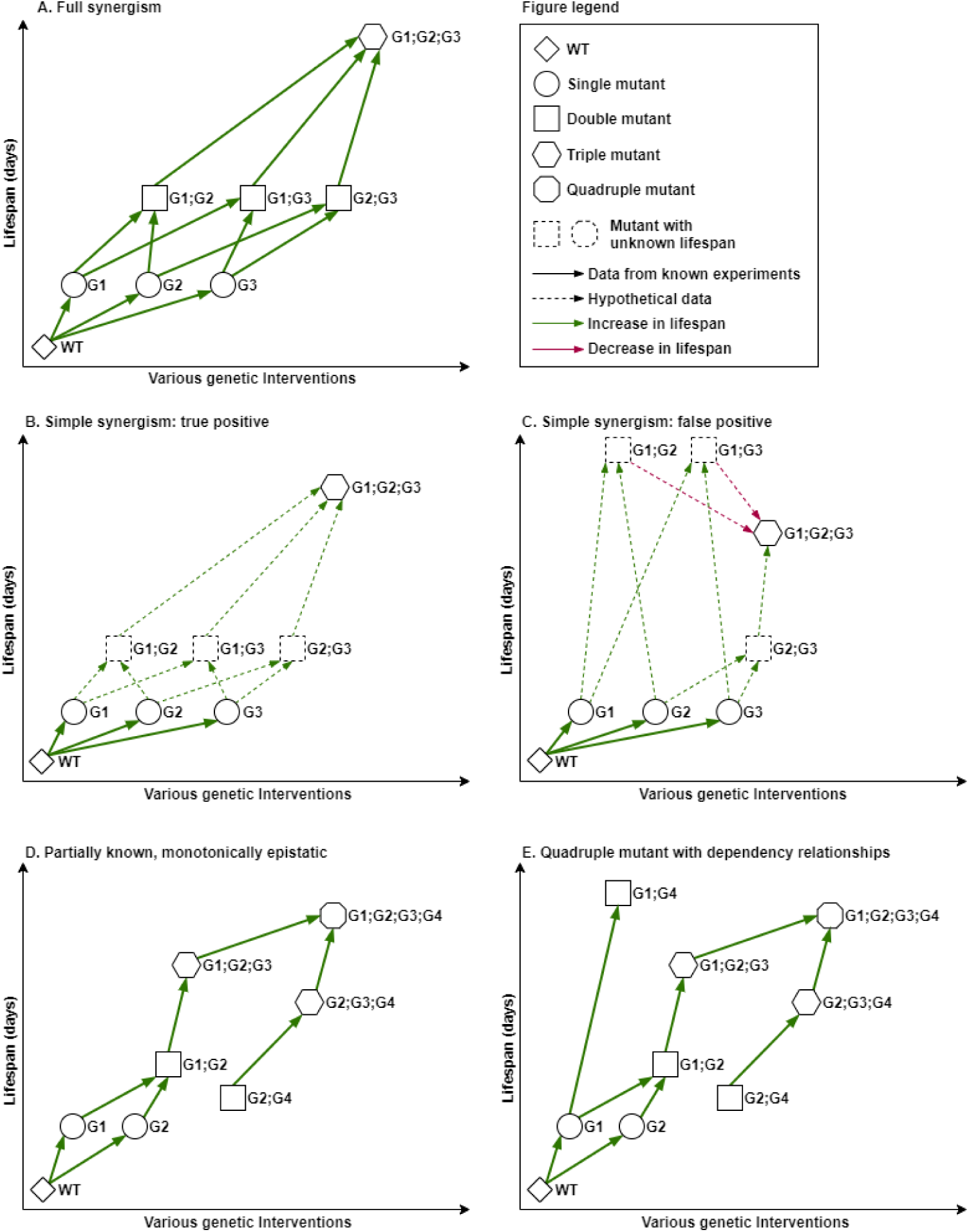
Synergism and epistatic monotony. **A.** Example of “full” synergism shown for a 3-mutant. **B.** True positive example of “simple” synergism approximation, in which the synergistic combination is correctly inferred based on the available data. The missing data (dotted nodes and edges) would not contradict the prediction. **C**. False positive example for the “simple” synergism approximation. The example shows an “incorrectly inferred” synergistic combination which may be disproved later by data that is currently unknown (dotted nodes G1;G2 and G1;G3 are even longer lived than G1;G2;G3). In this particular case, the dependency between G2 and G3 is masked by the missing lifespan values of intermediary mutants. Interventions in G2 and G3 significantly extend the lifespan of G1 separately, however performing both interventions is detrimental to the G1 mutant (even though G2 and G3 are additive or even synergistic when performed in the wild type). **D**. Example of an observed positive monotony, in the case of a partially known combination of 4 genes. Each additional intervention seems to have an added value towards lifespan extension. **E.** A quadruple mutant with dependency relationships. The 4-mutant is long lived, however the effect can be obtained by modulating G1, G4 only. Moreover, the effect of G1;G4 is even higher - suggesting that this combination is dependent on G2 and G3 not being affected. Of note, this occurs, regardless of G1 and G2 having an additive or synergistic relationship at lower levels (2-mutant G1; G2).

One practical issue still remaining is that while the above definitions are highly accurate, in practice, due to costs and time requirements, it is extremely unlikely for studies to provide lifespan curves for all the intermediary mutants. This is especially problematic when assaying higher level n-mutants. Moreover, even considering future progress in the development of automated screening platforms, the situation will change only slightly, as the combinatorial increase in the number of intermediary mutants is exponential. As such, it is our opinion that considering only “full” synergism limits the scope of any analysis either to a very small number of mutants (or even to a theoretical discussion only). Instead, approximations of synergism, or more generally of epistasis, could be used to evaluate the additive effects of lifespan interventions. Two such solutions are proposed below.

#### Approximation of a “simple” synergism

One simplification of synergism assessment would be to compare the lifespan effect of the n-gene combination only to the sum of the individual effects, and this is, in fact, the most popular definition of synergism (i.e. “the epistasis of two or more genes producing a combined effect greater than the sum of their separate effects”). The main advantage of this method is that in order to evaluate synergism, lifespan values are required only for the single mutants and the final n-mutant (see Fig. 2B). However, the main disadvantage of this approximation is that a subgroup of genes with a strong synergism could easily mask the non-significant, or perhaps even negative contribution of other genes that would falsely be considered synergistic to the aforementioned subgroup.

Unfortunately, due to the non-exhaustive nature of lifespan assays in published studies, this is a highly prevalent situation. An example of incorrectly inferred “simple” synergism is presented in Fig. 2C. An even weaker definition of the “simple” synergism would be that the lifespan effect of the gene combination is greater than all the individual gene effects, but not necessarily than the sum. This definition would produce an even higher number of false positives, so we do not consider it a viable option (for 2-mutants, this category is easily determined and was termed “almost” additive).

#### Partially known, monotonically epistatic

N-mutants can be constructed by performing a series of single gene interventions, in sequential order, with multiple paths. For example, in Fig. 2A, six paths can be observed, while in general, the number of paths is N = n!. If so, another more feasible approximation to evaluate epistasis between n genes could be to verify the monotonically increase or decrease in lifespan, however only along the paths that have been experimentally assayed. In this case, for a known path, if each genetic intervention k increases the lifespan effect of the previously combined k-1 interventions, then the path is monotonically positive (Fig. 2D). Similarly, a monotonically negative trend can be defined. Moreover, if at each step, the increase (or decrease) is equal or greater than the lifespan effect of the individual k intervention, then the n genes are synergistic in the “simple” sense (see previous approximation). It should be noted that this approximation does not allow for classifying the epistasis effect as synergistic, additive or dependent, since paths are missing and lifespan assay has not been performed for all mutants (see Fig. 2E for an example similar to that in Fig. 2D, but with one additional lifespan value). Nevertheless, when data is missing, this method can estimate the monotony of epistasis and provide a partially known trend.

To put into perspective the feasibility of implementing each of the above definitions, out of the total of 516 mutants constructed with 3 or more genetic interventions, and considering the available survival curves for intermediary mutants, “full” synergism can be evaluated for 21 mutants, “simple” synergism can be approximated for 29 mutants, and partially known monotonic epistasis can be estimated for 207 mutants (mutants having at least one path, starting with WT). Additionally, in the case of 2-mutants, full synergism can be evaluated for 1590 mutants and epistasis monotony for 1889 mutants. Currently, SynergyAge automatically annotates gene combinations based on the “full” synergism definition when all mutant data is available (Table 2), and computes the partially known monotony for all incomplete cases.

It should be stressed that all definitions above refer to data generated in the same experiment. One additional option that should be discussed is inferring synergism by including data from multiple experiments. For example, the lack of a mutant or wild-type in a certain experiment could be approximated by averaging those found in other studies (under the same conditions). This option, however, seems extremely unreliable as lifespan values vary significantly among different experiments.

For example, the range of recorded lifespans in our database for wild type strains of *C. elegans* under various conditions varies between 17.2 and 34.5 days at 15 degrees (avg: 23.66 days; SD: 4.36 days), and between 10.95 and 26 days at 20 degrees (avg: 18.8 days; SD 2.67 days) (for full histograms of WT lifespan at various temperatures, see http://www.synergyage.info/details/wildtype/). This was somewhat expected and is in line with literature reports as significant differences between the lifespan of controls is reported even in mice ^20^. Similarly, if we consider the lifespan variation for one of the most well-known long-lived mutants, *daf-2(e1370)* worms live between 32.3 and 48.7 days at 15 degrees (avg: 37.7 days; SD: 4.52 days), and between 21.86 and 60.8 days at 20 degrees (avg: 39 days; SD: 7.98 days). While SynergyAge has an inclusive approach and all data (including incomplete) is being collected and hosted, at this time data from multiple experiments is not used for epistasis inferences due to the large variations observed in different labs and experimental conditions.

Genes previously not associated with longevity that act as enhancers. The curation process for SynergyAge does not implicitly exclude experiments in which some of the assayed mutant populations display only small effects on longevity (e.g. lifespan increase < 10%), as long as an incremental increase/decrease is observed in combination with other genes. This type of inclusiveness might allow for identifying novel molecular players with potentially important roles in lifespan determination. Potentiators/enhancer genes could be synergistic with some of the already known LAGs, while not necessarily being LAGs themselves. Such examples, include but are not limited to: clk-1(e2519) or rsks-1(ok1255) enhancing the lifespan effect of daf-2(e1370), pep-2(lg601) enhancing the effect of let-363(RNAi), and asm-3(ok1744) potentiating age-1(mg305)’s effect.

Even more surprisingly, combining interventions with opposite lifespan effects can result in a shorter- or longer-lived double mutant, although such cases are somewhat rare (about 8.7% of the double mutants for which full synergism can be computed; 50 such long-lived mutants and 42 short-lived ones). Particularly interesting, among these are interventions that by themselves decrease lifespan and yet enhance the longevity effect of another lifespan-extending gene intervention (Supplementary Table S1). For example, the double mutant *daf-2(e1370);pep-2(lg601)* has an increase in lifespan compared to wild type, twice greater than the single mutant *daf-2(e1370)*, even though *pep-2(lg601)* is a short-lived mutant.

## TECHNICAL VALIDATION

The SynergyAge data was validated at three levels: 1) during the curation step, 2) before the insertion to the database, and 3) after database deployment, manually investigating the outliers and random entries in the final data. With regards to the data source, only papers with experimentally validated results were selected for curation, hence the data had also a layer of pre-validation. In addition, the survival curves presented in the included papers were manually verified, as described in the methods section. When importing the data, automatic checks were performed to ensure there are no missing values in the text files or a wrong format in the data. Following the data import, lifespan distributions for each mutant were plotted to manually investigate the outliers at different temperatures.

## USAGE NOTES

The SynergyAge interface is an easy-to-use web-platform, in which data can be viewed either by browsing through all the mutant strains of an organism or by using the search function. The text search query can contain any number of genes, separated by comma or semicolon. Results are paginated when browsing, and are displayed grouped in four categories when searching (Fig. 3):

1. mutants with genetic interventions in all of the searched genes (if such a mutant exists), a category with results useful especially when searching for a specific n-mutant;
2. 1-mutants, generated by single genetic modifications in any of the genes from the search list, thus offering the option to “batch”-search for multiple 1-mutants and also to use partial-name search to identify genes present in the database;
3. n-mutants constructed based on several genetic interventions corresponding to a subset of the searched genes, a result section that can be used to identify lower degree mutants (for example to identify the 2-mutants that are related to a certain 3-mutant);
4. combinations that include at least *one* or *all* (depending on a user-selectable filter) of the searched genes but also other genes that were not searched for, an option that can be used for an expansive search starting from a simple model and exploring towards more complex ones.

**Fig. 3.**
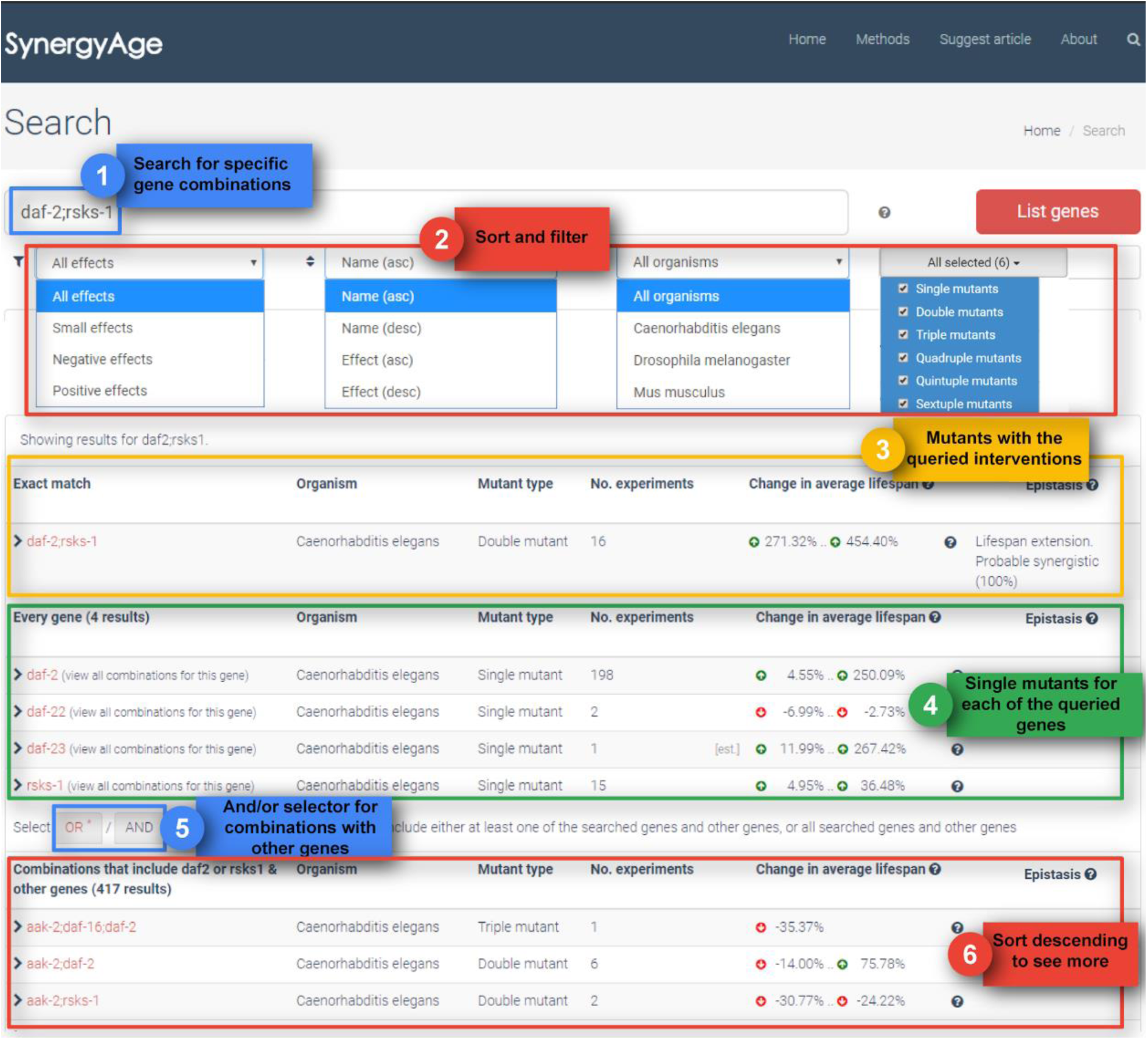
Usage example of the search interface in SynergyAge. Results are displayed for the daf-2/rsks-1 query. Depicted are the sort and filter options, and the returned list of results group into categories. The y coordinate of a node is proportional to their lifespan. The position of mutants on the x-axis is ascendingly ordered by the number of genetic interventions.

In addition, during browsing and searching, several other sorting and filtering options are also available: filtering by lifespan effect (positive, negative of small effects), filtering by the number of mutations (the n-degree of the mutant), and sorting by the name of the mutant or by the change in average lifespan. The table-form results include the number of experiments from which the data was extracted, an average of the lifespan changes obtained with the n-mutant, and the estimated epistasis relationship between the modulated genes. For gene combinations annotated with more than one epistatic relationship due to multiple literature results, the “Epistasis” column summarizes all the possible relationships for a particular combination (Fig. 3). Conversely, if the data in the database is incomplete (data for some of the “intermediary” mutants not being available), a “Not enough data to assess” message is provided.

A special effort has been dedicated to the development of an interactive module that allows the user to visually surf through a network of mutants for each species (Fig. 4A). In the networks, each node represents a lifespan model (wild type or n-mutant), while edges indicate the relationship between models, emphasizing for a certain intervention the background in which it was done (the n-mutant) and the mutant with one more modified gene (the n+1 mutant) that was obtained (e.g.: wild type >=> 1-mutant, 1-mutant >=> 2-mutant, etc).

**Fig. 4.**
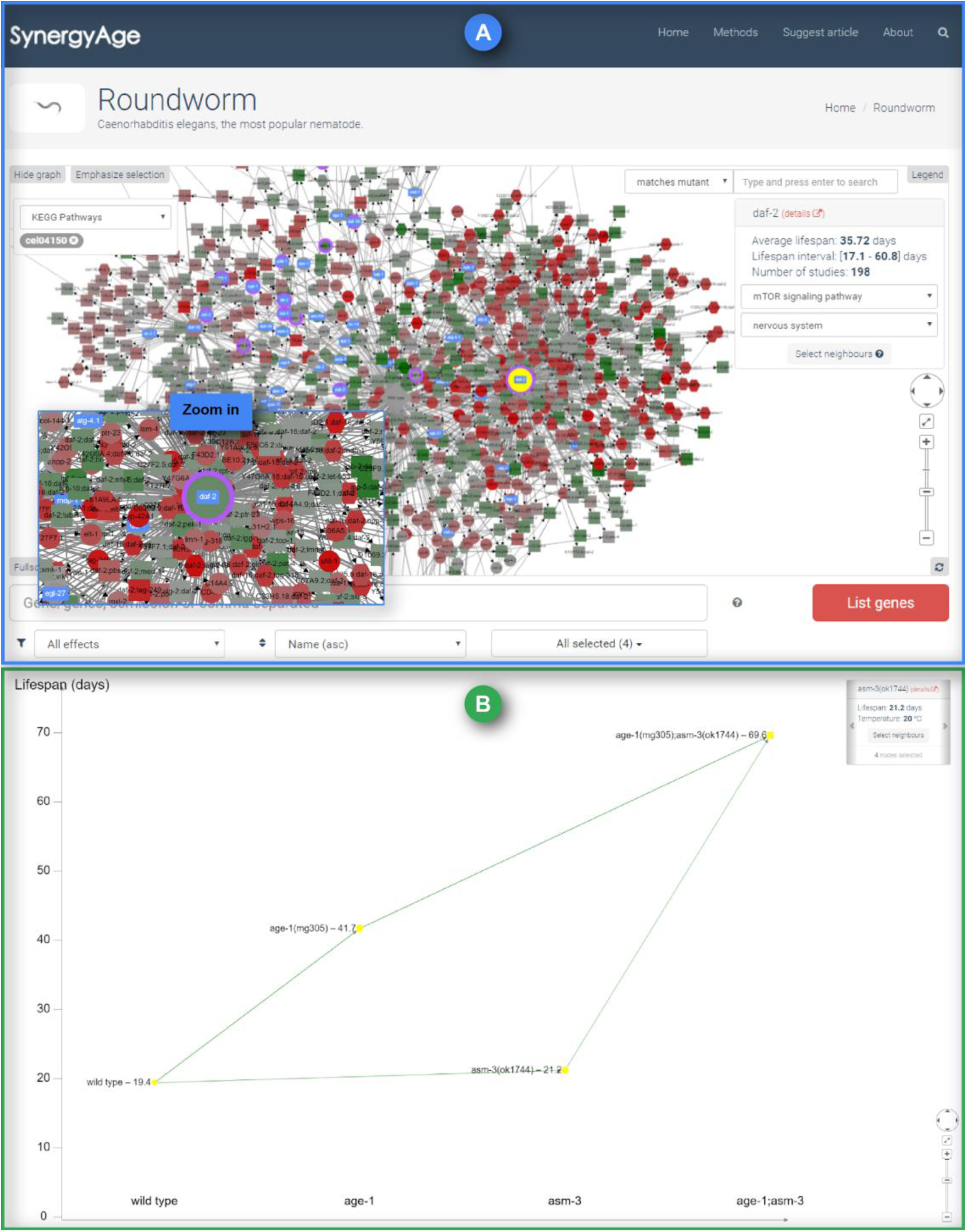
The network visualisation module. **A.** The graph representation of all *C.elegans* mutants recorded in the database. Mutant models with more data entries, due to multiple lifespan assay studies, can be easily identified by larger node sizes. Colors are used to help identify the direction and the intensity of the lifespan change for each genetic intervention. Green indicates long-lived mutants, red points to short-lived ones. The color intensity of the node is proportional to the change in average lifespan from all experiments pertaining to a node in the database, relative to the average wild type in the database. Purple border and/or blue overlay can be used to mark all single gene mutants in the network related to a selected pathway/tissue. **B.** Example of an individual entry page for the gene combination age-1/asm-3. Green transitions indicate additional interventions that result in longer-lived mutants, red transitions point to additional interventions that are more detrimental - shorter-lived mutants.

In the network module, node selection can be done either directly (mouse click/shift mouse click or shift+selection rectangle) or using the module’s search text field. If using the in-network search field, the type of search can be either: 1) exactly matching a mutant name (“matches mutant”), or 2) a partial search (“contains gene”), selecting mutants that contain the gene of interest. Selected nodes in the network are highlighted in yellow and will be included in the carousel panel at the top right corner of the module. For all selected nodes, the carousel panel provides links to each mutant’s page and basic lifespan statistics (average lifespan, lifespan interval and number of studies from which data was gathered). It is worth mentioning that a fast option to select the neighbors of the currently displayed node in the panel also exists (neighbors will be added to the current selection). In order to allow the user to focus on a certain part of the network, the “Emphasize selection” button can fade out the non-selected nodes.

For 1-mutants, the network also integrates external data, from KEGG for pathways and from model-specific databases (e.g. Wormbase) for gene expression in tissues. This way, the carousel panel can be used to visually mark all the single mutants (respectively the genes) in the network that are associated with the same pathway or expressed in the same tissue as the currently selected gene.

For each n-mutant, the website contains a page with detailed information on individual experiments. On this page, the information is grouped based on the type of genetic interventions, and for each type, lifespan data together with experimental conditions from each experiment are included. Short summaries and a quick link to the published work are also included, allowing users to investigate the reports in more detail. Complementing these tables, the page also includes a graphical representation of all the reported effects. Lifespan values are included in this graph for each experiment separately (Fig. 4B). Nodes represent individual cohorts and edges between nodes are drawn only between animal populations from the same experiment.

To visually evaluate the epistatic effect of a gene combination, users can click on an n-mutant of interest and then use either “Select neighbors” from the carousel panel (obtaining a list of n-1 and n+1 mutants), or using the button “Select genes with PubMed …” (obtaining all the mutants from the same paper).

## CODE AVAILABILITY

The source for the website code and database schema are available from a public github repository (https://github.com/gerontomics/synergyage).

## DATA AVAILABILITY

The SynergyAge database is publicly available at www.synergyage.info. The data is made available under the permissive Creative Commons license and may be freely used in other analyses. Additionally, the SynergyAge database includes a download option which allows for an export of the data as a JSON file.

## Contributions

This study was carried out by the research groups led by RT, VEF and JPM. RT and GB conceptually designed the project, and RT supervised the entire implementation. Data collection and processing were carried out by GB, DT, LS and DB. Automatization of importing, bioinformatics and statistical analyses were carried mainly by GB. The website, networks and database were built by CFI and GB. LGG supported the project with server deployment, administration and security. Data validation and website testing were done by RT, GB, DT and LS. All authors have participated in the writing of the manuscript, with major contributions from GB and DT. Additionally, RT, JPM and VEF have ensured the quality of the manuscript and its scientific correctness. Figures were created by DT and GB. All authors have read and approved the final manuscript.

## Supplementary materials

**Table S1.** The enhancers with the opposite lifespan effects.

